# Improving the diagnostic yield of exome-sequencing, by predicting gene-phenotype associations using large-scale gene expression analysis

**DOI:** 10.1101/375766

**Authors:** Patrick Deelen, Sipko van Dam, Johanna C. Herkert, Juha M. Karjalainen, Harm Brugge, Kristin M. Abbott, Cleo C. van Diemen, Paul A. van der Zwaag, Erica H. Gerkes, Pytrik Folkertsma, Tessa Gillett, K. Joeri van der Velde, Roan Kanninga, Peter C. van den Akker, Sabrina Z. Jan, Edgar T. Hoorntje, Wouter P. te Rijdt, Yvonne J. Vos, Jan D.H. Jongbloed, Conny M.A. van Ravenswaaij-Arts, Richard Sinke, Birgit Sikkema-Raddatz, Wilhelmina S. Kerstjens-Frederikse, Morris A. Swertz, Lude Franke

## Abstract

Clinical interpretation of exome and genome sequencing data remains challenging and time consuming, with many variants with unknown effects found in genes with unknown functions. Automated prioritization of these variants can improve the speed of current diagnostics and identify previously unknown disease genes. Here, we used 31,499 RNA-seq samples to predict the phenotypic consequences of variants in genes. We developed GeneNetwork Assisted Diagnostic Optimization (GADO), a tool that uses these predictions in combination with a patient’s phenotype, denoted using HPO terms, to prioritize identified variants and ease interpretation. GADO is unique because it does not rely on existing knowledge of a gene and can therefore prioritize variants missed by tools that rely on existing annotations or pathway membership. In a validation trial on patients with a known genetic diagnosis, GADO prioritized the causative gene within the top 3 for 41% of the cases. Applying GADO to a cohort of 38 patients without genetic diagnosis, yielded new candidate genes for seven cases. Our results highlight the added value of GADO (www.genenetwork.nl) for increasing diagnostic yield and for implicating previously unknown disease-causing genes.

## Introduction

With the increasing use of whole-exome sequencing (WES) and whole-genome sequencing (WGS) to diagnose patients with a suspected genetic disorder, diagnostic yield is steadily increasing [1]. Although our knowledge of the genetic basis of Mendelian diseases has improved considerably, the underlying cause remains elusive for a substantial proportion of cases. The diagnostic yield of genome sequencing varies from 8% to 70% depending on the patient’s phenotype and the extent of genetic testing [2]. Sequencing all ∼20,000 protein-coding genes by WES and entire genomes by WGS usually increases sensitivity but decreases specificity: it results in off-target noise and reveals many variants of uncertain clinical significance. In a study by Yang *et al.*, proband-only WES identified approximately 875 variants in each patient, even after removing low quality variants [3].

One strategy to manage the list of genetic variants is to perform trio analysis of samples from the proband and both of his or her biological parents to ascertain, for instance, whether a variant has *de novo* status [4]. Another strategy is to limit the analyses to a gene panel of Online Mendelian Inheritance in Men (OMIM) disease-annotated genes [5] or genes known to be directly related to the patient’s phenotype. However, determining the actual disease-causing variant requires further variant filtering based on information about its predicted functional consequence, population frequency data, conservation, disease-specific databases (such as the Human Gene Mutation Database [6]), literature, and segregation analysis [7].

Several tools have been developed that aid in variant filtering and prioritization [8,9]. Annotation tools, such as VEP [10] and GAVIN [9], offer additional functionality that allows variants to be filtered according to their population frequency and variant class. Other tools use phenotype descriptions to rank potential candidates genes [11]. The phenotypes are typically described in a structured manner, e.g. using Human Phenotype Ontology (HPO) terms [12]. AMELIE (Automatic Mendelian Literature Evaluation), for example, prioritizes candidate genes by their likelihood of causing the patient’s phenotype based on automated literature analysis [13]. However, this focus on what is known may inadvertently filter out variants in potential novel disease genes. Alternatively, the causative gene defect could be missed if a patient’s phenotype differs from the features previously reported to be associated to a disease gene. Tools like Exomiser can identify novel human disease genes, as it prioritizes variants based on semantic phenotypic similarity between a patient’s phenotype described by HPO terms and HPO-annotated diseases, Mammalian Phenotype Ontology (MPO)-annotated mouse and Zebrafish Phenotype Ontology (ZPO)-annotated fish models associated with each exomic candidate and/or its neighbors in an interaction network [14]. However, most available algorithms are based on existing knowledge on human disease genes, their orthologues in animal models, or well-described biological pathways (for a detailed review see [11]).

To overcome this, we hypothesized that co-regulation of expression data could be used to prioritize variants, including those in less well studied genes. We assumed that if a gene or a gene set is known to cause a specific disease or disease symptom, these genes will often have similar molecular functions or be involved in the same biological process or pathway. We reasoned that variants in genes with yet unknown function that are involved in the same biological pathway or co-regulated with known disease genes likely result in the same phenotype. In order to identify groups of genes with a related biological function, we used an expansive compendium of 31,499 RNA-sequencing (RNA-seq) gene expression samples to predict functions for genes with high accuracy.

We then developed a user-friendly tool that can prioritize variants in known *and* unknown genes based on our functional predictions, which we designated GeneNetwork Assisted Diagnostic Optimization (GADO). GADO ranks variants based on gene co-regulation in publicly available expression data of a wide range of tissues and cell types using HPO terms to describe a patient’s phenotype. To validate our prioritization method, we tested how well our method predicts disease-causing genes based on features described for each of the genes in the OMIM database. We then used exome sequencing data of patients with a known genetic diagnosis to benchmark GADO. Finally, we applied our methodology to previously inconclusive WES data and identified several genes that contain variants that likely explain the phenotype of the respective patients. Thus, we show that our methodology is successful in identifying variants in novel, potentially relevant genes explaining the patient’s phenotype.

## Results

### Gene prioritization using GADO

We have developed GADO to perform gene prioritizations using the phenotypes observed in patients denoted as HPO terms [15]. In combination with a list of candidate genes (i.e. genes harboring rare and possibly damaging variants), this results in a ranked list of genes with the most likely candidate genes on top (**Figure 1a**). The gene prioritizations are based on the predicted involvement of the candidate genes for the specified set of HPO terms. These predictions are made by analyzing public RNA-seq data from 31,499 samples (**Figure** 1b), resulting in a gene prediction score for each HPO term. These predictions are solely based on co-regulation of genes annotated to a certain HPO term with other genes. This makes it possible to also prioritize genes that currently lack any biological annotation.

**Figure 1:**
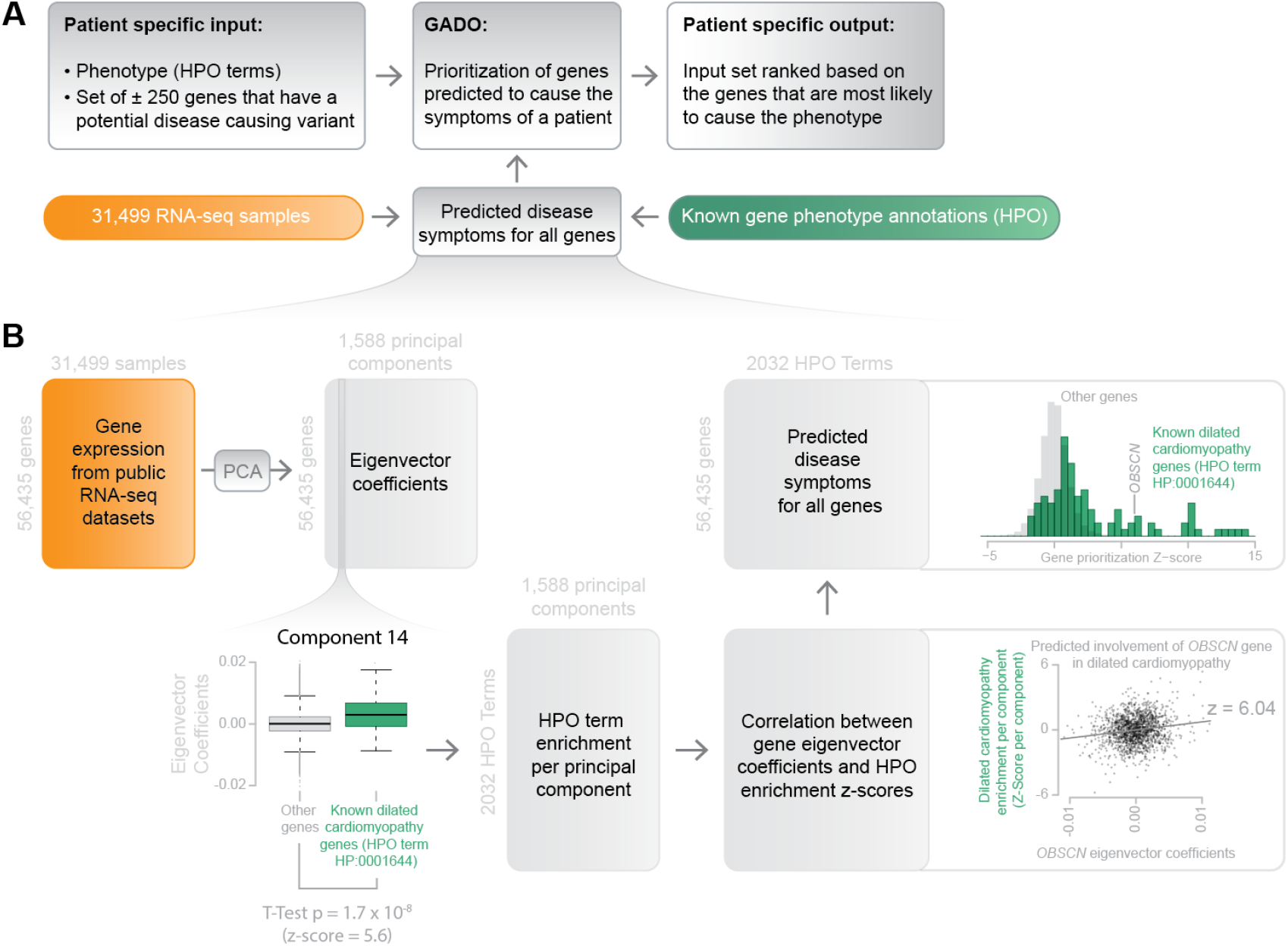
Schematic overview of GADO. (a) Per patient, GADO requires a set of phenotypic features and a list of candidate genes (i.e. genes harboring rare alleles that are predicted to be pathogenic) as input. It then ascertains whether genes have been predicted to cause these features, and which ones are present in the set of candidate genes that has been provided as input. The predicted HPO phenotypes are based on the co-regulation of genes with sets of genes that are already known to be associated with that phenotype. (b) Overview of how disease symptoms are predicted using gene expression data from 31,499 human RNA-seq samples. A principal component analysis on the co-expression matrix results in the identification of 1,588 significant principal components. For each HPO term we investigate every component: per component we test whether there is a significant difference between eigenvector coefficients of genes known to cause a specific phenotype and a background set of genes. This results in a matrix that indicates which principal components are informative for every HPO term. By correlating this matrix to the eigenvector coefficients of every individual gene, it is possible to infer the likely HPO disease phenotype term that would be the result of a pathogenic variant in that gene.

### Public RNA-seq data acquisition and quality control

To predict functions of genes and HPO term associations, we downloaded all human RNA-seq samples publicly available in the European Nucleotide Archive (accessed June 30, 2016) (supplementary table 1) [16]. We quantified gene-expression using Kallisto [17] and removed samples for which a limited number of reads are mapped. We used a principal component analysis (PCA) on the correlation matrix to remove low quality samples and samples that were annotated as RNA-seq but turned out to be DNA-seq. In the end, we included 31,499 samples and quantified gene expression levels for 56,435 genes (of which 22,375 are protein-coding).

Although these samples are generated in many different laboratories, we previously observed that, after having corrected for technical biases, it is possible to integrate these samples into a single expression dataset [18]. We validated that this is also true for our new dataset by visualizing the data using t-Distributed Stochastic Neighbor Embedding (t-SNE). We labeled the samples based on cell-type or tissue and we observed that samples cluster together based on cell-type or tissue origin (**Figure 2a**). Technical biases, such as whether single-end or paired-end sequencing had been used, did not lead to erroneous clusters, which suggests that this heterogeneous dataset can be used to ascertain co-regulation between genes and can thus serve as the basis for predicting the functions of genes.

**Figure 2:**
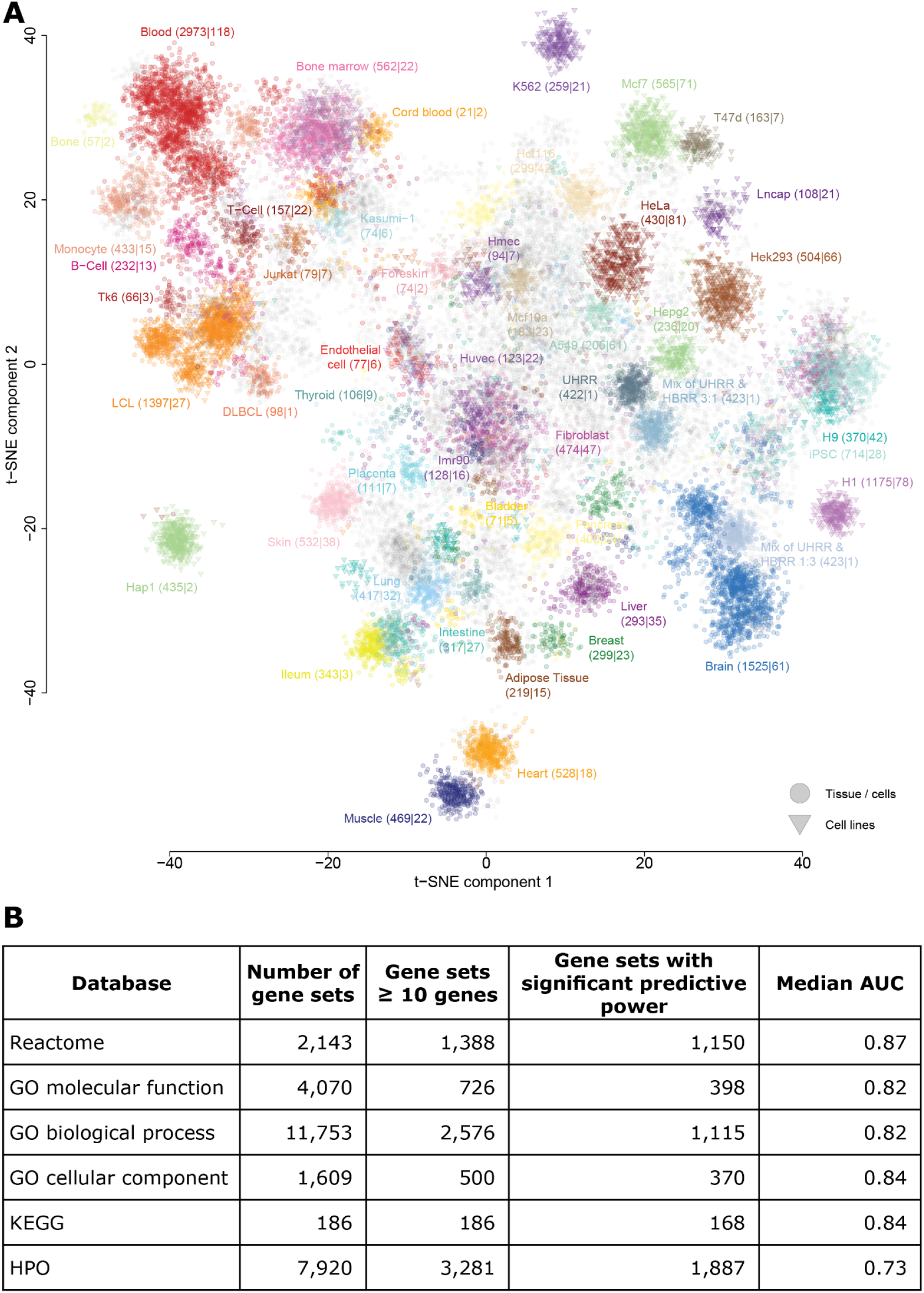
A compendium of gene expression profiles that can be used for gene function prediction. (a) 31,499 RNA-seq samples derived from many different studies show coherent clustering after correcting for technical biases. Generally, samples originating from the same tissue, cell-type or cell-line cluster together. The two axes denote the first t-SNE components. (b) Gene co-expression information of 31,499 samples is used to predict gene functions. We show the prediction accuracy for gene sets from different databases. AUC, Area Under the Curve, GO, Gene Ontology, HPO, Human Phenotype Ontology.

### Prediction of gene HPO associations and gene functions

To predict HPO term associations and putative gene functions using co-regulation (**Figure 1b**), we used a method that we had previously developed and applied to public expression microarrays [19]. Since these microarrays only cover a subset of the protein-coding genes (n = 14,510), we decided to use public RNA-seq data instead. This allows for more accurate quantification of lower expressed genes and the expression quantification of many more genes, including a large number of non-protein-coding genes. [20].

We applied this prediction methodology [19] to the HPO gene sets and also to Reactome [21], KEGG pathways [22], Gene Ontology (GO) molecular function, GO biological process and GO cellular component [23] gene sets. For 5,088 of the 8,657 gene sets (59%) with at least 10 genes annotated, the gene function predictions had significant predictive power (see materials and methods). For the 8,657 gene sets with at least 10 genes annotated, the median predictive power, denoted as Area Under the Curve (AUC), ranged between 0.73 (HPO) to 0.87 (Reactome) (**Figure 2b**).

### Prioritization of known disease genes using the annotated HPO terms

Once we had calculated the prediction scores of HPO disease phenotypes, we leveraged these scores to prioritize genes found by sequencing the DNA of a patient. For each individual HPO term–gene combination, we calculated a prediction z-score that can be used to rank genes. In practice, however, patients often present with not one feature but a combination of multiple features. Therefore, we combined the z-scores for each HPO term [24] to generate an overall z-score that explains the full spectrum of features in a patient. GADO uses these combined z-scores to prioritize the candidate genes: the higher the combined z-score for a gene, the more likely it explains the patient’s phenotype.

Because many HPO terms have fewer than 10 genes annotated, and since we were unable to make significant predictions for some HPO terms, certain HPO terms are not suitable to use for gene prioritization. We solved this problem by taking advantage of the way HPO terms are structured. Each term has at least one parent HPO term that describes a more generic phenotype and thus has also more genes assigned to it. Therefore, if an HPO term cannot be used, GADO will make suggestions for suitable parental terms (supplementary figure 1).

To benchmark our prioritization method, we used the OMIM database [5]. We tested how well our method was able to retrospectively rank disease-causing genes listed in OMIM based on the annotated symptoms of these diseases. We took each OMIM disease gene (n = 3,382) and used the associated disease features (15 per gene on average) as input for GADO. What we found was that for 49% of the diseases GADO ranks the causative gene in the top 5% (**Figure 3a, b**). Moreover, we observed a statistically significant difference between the performance of GADO on true gene-phenotype combinations and its performance using a random permutation of gene-phenotype combinations (p-value = 2.16 × 10^-532^).

**Figure 3:**
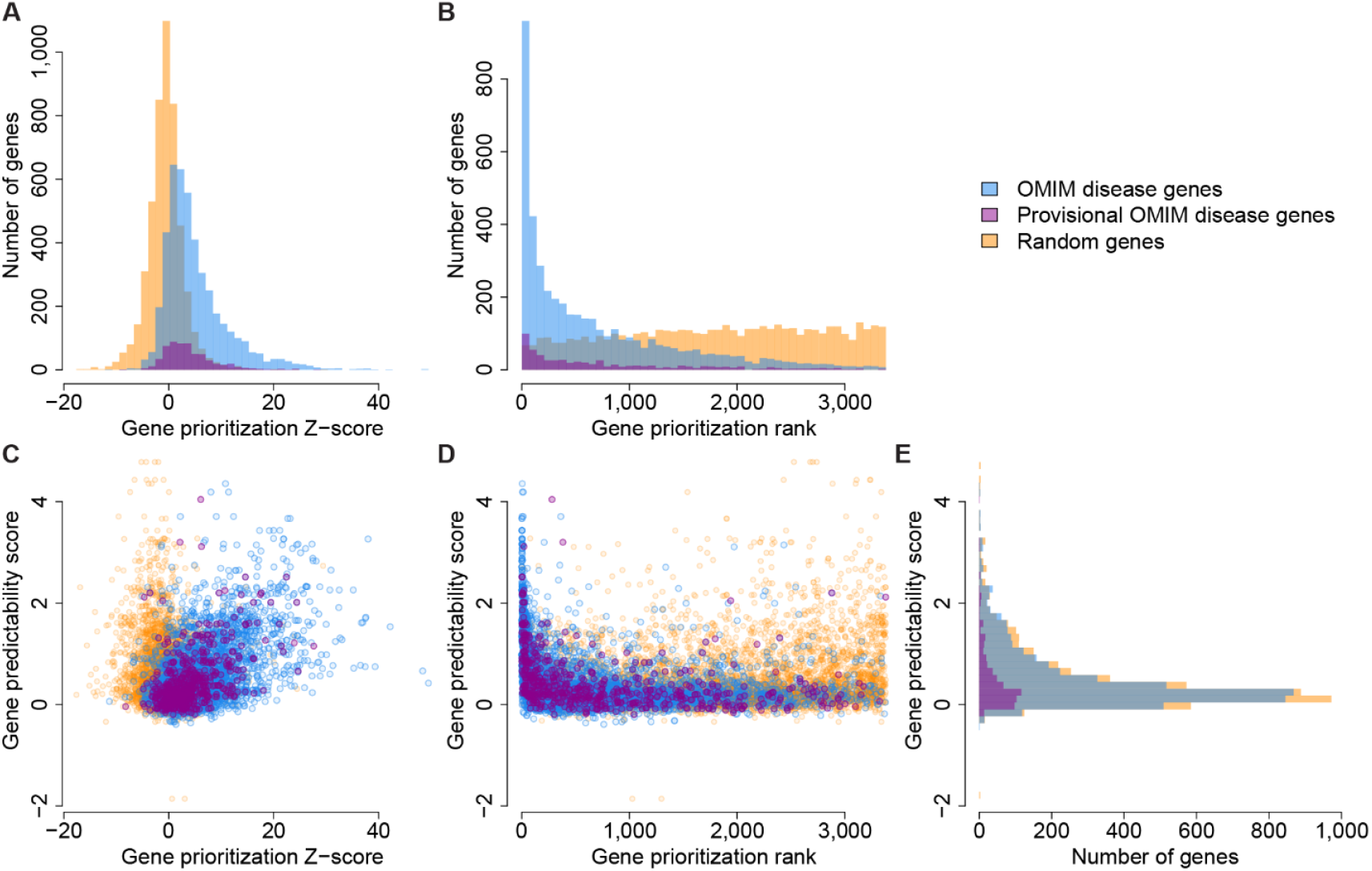
Performance of disease gene prioritization compared to random permutation. (a) OMIM disease genes and provisional disease genes have significantly stronger z-scores compared to permuted disease genes (T-test p-values: 2.16×10^-532^ & 5.38×10^-80^, respectively). We also observe that the predictions of the provisional OMIM genes are, on average, weaker than the other OMIM disease genes (T-test p-value: 1.89×10^-7^). (b) Ranking the disease based on z-scores shows GADO’s ability to prioritize the causative gene for a disease among all OMIM genes. For 49% of the disorders the causative gene is ranked in the top 5%. (c) We observe a clear relation between the prioritization z-scores and the gene predictability scores (Pearson r = 0.54). We don’t observe this relation in the permuted results. (d) GeneNetwork performs best for genes with high predictability scores. (e) The different groups have similar distributions of gene predictability scores.

### Gene predictability scores explains performance differences between genes

For some combinations of genes and HPO terms listed in OMIM, GADO could not establish the gene-phenotype combination (**Figure 3**). For example, variants in *SLC6A3* are known to cause infantile Parkinsonism-dystonia (MIM 613135) [25–27], but GADO was unable predict the annotated HPO terms related to the Parkinsonism-dystonia for this gene. This may, however, be due to very low expression levels of *SLC6A3* in most tissues except specific brain regions [28].

To better understand why we can’t predict HPO terms for all genes, we used the Reactome, GO and KEGG prediction scores. Jointly these databases comprise thousands of gene sets. Since these databases describe such a wide range of biology, we assumed that if a gene does not show any prediction signal for any gene set in these databases, gene co-expression is probably not informative for this gene. To quantify this, we calculated, per gene, the average skewness of the z-score distribution of the Reactome, GO and KEGG gene sets. From this we were able to derive a ‘gene predictability score’ for every gene that is independent of whether this gene is already known to play a role in any a disease or pathway (**Figure 3c, d, e**). We then ascertained whether these ‘gene predictability scores’ are correlated with the prediction z-score of the OMIM diseases, and found a strong correlation (Pearson r = 0.54, p-value = 1.14 × 10^-332^) between the gene predictability scores and GADO’s ability to identify a known disease gene (**Figure 3c**).

To investigate why some genes have a high ‘gene predictability score’ but low prediction performance, we scored a set of genes known to cause cardiomyopathy (CM) for the amount of literature evidence that these genes cause CM. We found several genes for which the prediction score for the CM phenotype is lower than expected based on the gene predictability scores (supplementary figure 2a). Pathogenic variants in the *TTR* gene implicated in hereditary amyloidosis (MIM 105210) [29], for instance, cause accumulation of the transthyretin protein in different organ systems, including the heart, resulting in CM. However, this gene is primarily expressed in the liver. Therefore, its disease mechanism is different from other mechanisms resulting in CM, as many inherited CMs are caused by deleterious variants in genes highly expressed in the heart and directly affecting the function of the cardiac sarcomere. Therefore, the phenotypic function prediction for this gene may be worse than we would expect based on the predictability score. We performed a similar analysis using the HPO term ‘dilated cardiomyopathy’ and observed a low prediction performance for the *TMPO* gene, despite a high gene predictability score (supplementary figure 2b). Previously, this gene was reported to be related to dilated cardiomyopathy (DCM) and listed as such by OMIM. However, recent reclassification of the reported variants using the ExAC data revealed that the reported variant was far too common to be causative for DCM [30].

### Benchmarking GADO using solved cases with realistic phenotyping

Although *in silico* benchmarking demonstrated the potential of GADO, it used all annotated HPO terms for a disease. In practice, however, patients may only present with a limited number of the annotated features. To perform a validation that was a more realistic reflection of clinical practice, we used exome sequencing data of 83 patients with a known genetic diagnosis. We used their phenotypic features as listed in their medical records prior to the genetic diagnosis (supplementary table 2). On average, per patient, GADO yielded 56 possible disease-causing genes with variants that are rare and predicted to be deleterious. In 41% of the patients the actual causative gene was ranked in the top 3 and in 50% of the cases it was in the top 5 (mean rank 10) (**Figure 4a**).

**Figure 4:**
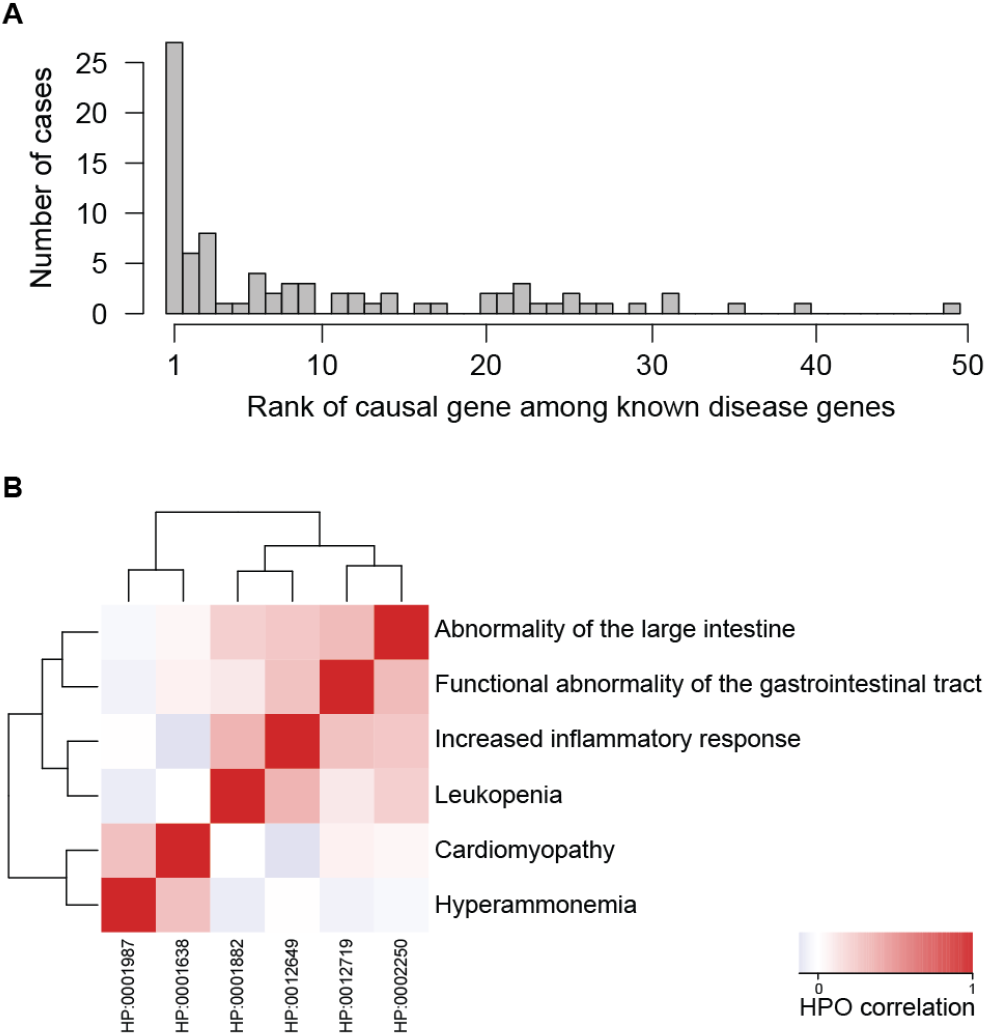
Performance of GeneNetwork on solved cases. (a) Rank of the known causative gene among the candidate disease causing variants. (b) Our cohort contained a case with two distinct conditions, and clustering showed the HPO terms of the same disease are closest to each other. Note, the HPO term “Inflammation of the large intestine” did not yield a significant prediction profile and therefore the parent terms “Abnormality of the large intestine”, “Increased inflammatory response” and “Functional abnormality of the gastrointestinal tract” where used for this case.

### Clustering of HPO terms

In addition to ranking potentially causative genes based on a patient’s phenotype, we observed that GADO can be used to cluster HPO terms based on the genes that are predicted to be associated to these HPO terms. This can help identify pairs of symptoms that often occur together, as well as symptoms that rarely co-occur, and we actually observed this for a patient suspected of having two different diseases. This patient is diagnosed with a glycogen storage disease, GSD type Ib, caused by compound heterozygous variants in *SLC37A4* (MIM 602671) and DCM that is probably caused by a truncating variant in *TTN* (MIM 188840). Clustering of the assigned HPO terms placed the phenotypic features related to GSD type Ib (‘leukopenia’ (HP:0001882) and ‘inflammation of the large intestine’ (HP:0002037)) together, while Cardiomyopathy (HP:0001638) was only weakly correlated to these specific features (**Figure 4b**).

### Reanalysis of previously unsolved cases

To assess GADO’s ability to discover new disease genes, we applied it to data from 38 patients who are suspected to have a Mendelian disease but who have not had a genetic diagnosis. All patients had undergone prior genetic testing (WES with analysis of a gene panel according to their phenotype, supplementary table 3). On average three genes had a z-score ≥ 5 (which we used as an arbitrary cut-off and that correspond to a p-value of 5.7 × 10^-7^) and were further assessed. In seven cases, we identified variants in genes not associated to a disease in OMIM or other databases, but for which we could find literature or for which we gained functional evidence implicating their disease relevance (**Table 1**). For example, we identified two cases with DCM with rare compound heterozygous variants in the *OBSCN* gene (MIM 608616) that are predicted to be damaging. In literature, inherited variant(s) in *OBSCN*, encoding obscurin, are associated with hypertrophic CM [31] and DCM [32]. Furthermore, obscurin is a known interaction partner of titin (TTN), a well-known DCM-related protein [31]. Another example came from a patient with ichthyotic peeling skin syndrome, which is caused by a damaging variant in *FLG2 (*MIM 616284). We recently published this case where we prioritized this gene using an alpha version of GADO [33].

**Table 1:**
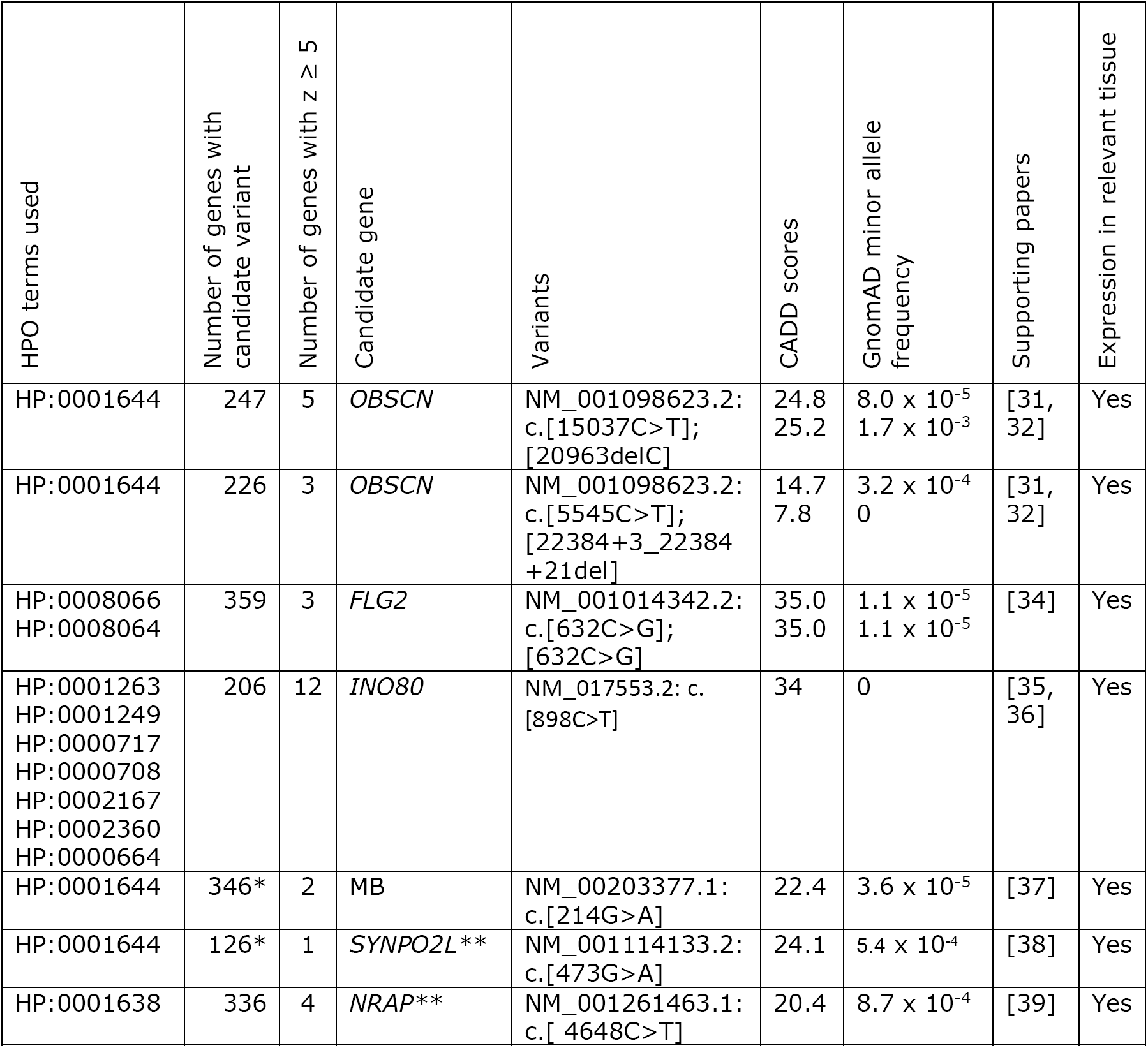
unsolved cases with new candidate genes. Out of the 38 unsolved patients investigated, we identified candidate genes in seven patients. For these genes we have found literature that indicates these genes fit the phenotype of these patients or for which we gained functional evidence implicating their disease relevance. *These variants where pre-filtered for family segregation. **The variants in these genes do not fully explain the phenotype but are likely contributing to the phenotype.

### www.genenetwork.nl

All analyses described in this paper can be performed using our online toolbox at www.genenetwork.nl. Users can perform gene prioritizations using GADO by providing a set of HPO terms and a list of candidate genes (**Figure 5a**). Per gene, it is also possible to download all prediction scores for the HPO terms and pathways. Our co-regulation scores between genes can be used for clustering. Furthermore, the predicted pathway and HPO annotations of genes can be used to perform function enrichment analysis (**Figure 5b**). We also support automated queries to our database.

**Figure 5:**
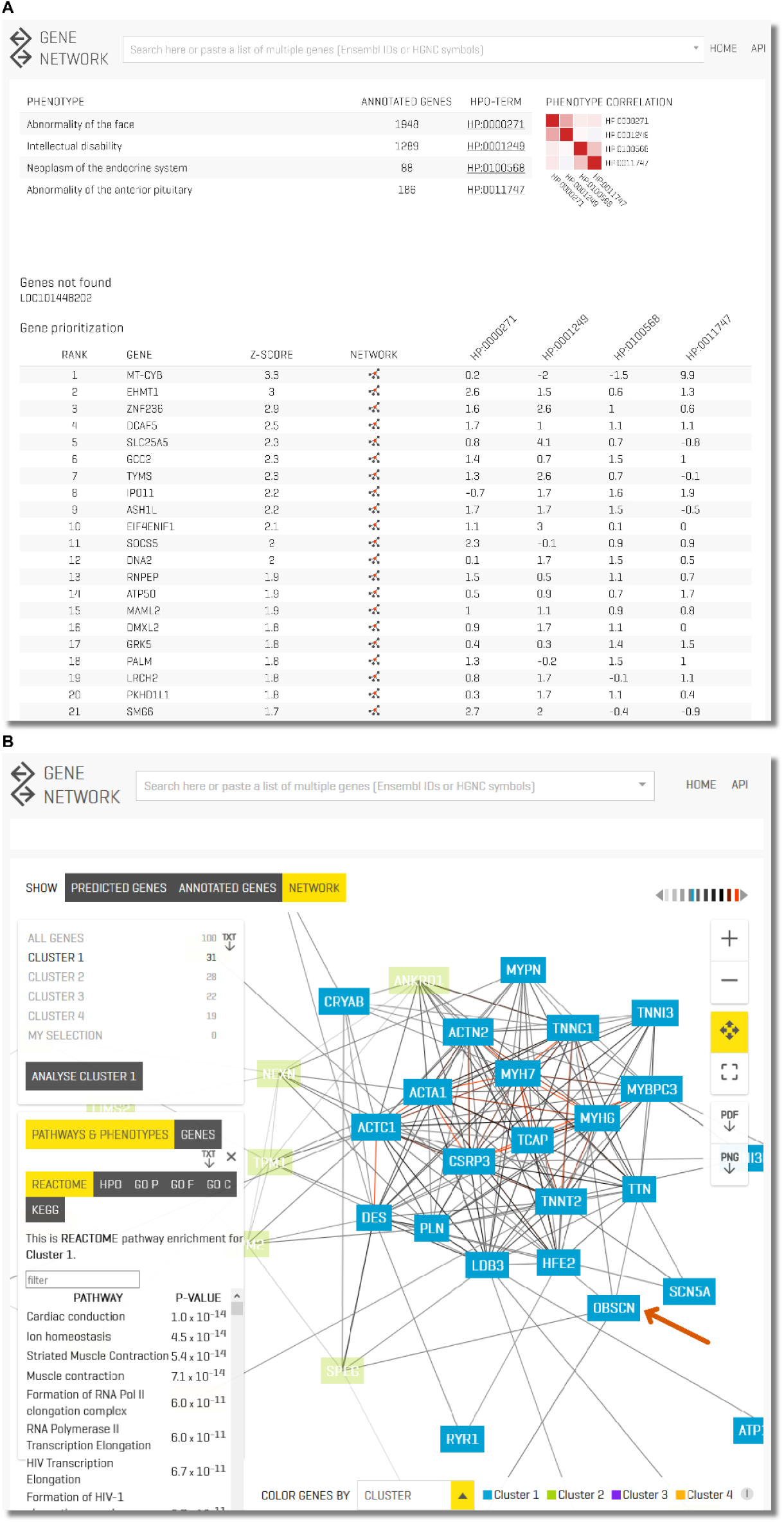
www.genenetwork.nl (a) Prioritization results of one of our previously solved cases. This patient was diagnosed with Kleefstra syndrome. The patient only showed a few of the phenotypic features associated with Kleefstra syndrome and additionally had a neoplasm of the pituitary (which is not associated with Kleefstra syndrome). Despite this limited overlap in phenotypic features, GADO was able to rank the causative gene (EHMT1) second. Here, we also show the value of the HPO clustering heatmap, the two terms related to the neoplasm cluster separately from the intellectual disability and the facial abnormalities that are associated to Kleefstra syndrome. (b) Clustering of a set of genes allowing function / HPO enrichment of all genes or specific enrichment of automatically defined sub clusters. Here we loaded all known DCM genes and OBSCN, and we focus on a sub-cluster of genes containing OBSCN (highlighted by the arrow). We see that it is strongly co-regulated with many of the known DCM genes. Pathway enrichment of this sub-cluster reveals that these genes are most strongly enriched for the muscle contraction Reactome pathway. DCM, Dilated Cardiomyopathy.

## Discussion

Prioritizing genes from WES or WGS data remains challenging. To meet this challenge, we developed GADO, a novel tool to prioritize genes based on the phenotypic features of a patient. Since the classification of variants is labor-intensive, prioritization of the most likely candidate variants saves time in the diagnostic process.

Importantly, GADO can also aid in the discovery of currently unknown disease genes. The main advantage of our methodology is that it does not rely on any prior knowledge about disease-gene annotations. Instead, we used predicted gene functions based on co-expression networks extracted from a large compendium of publicly available RNA-seq samples. RNA-seq has previously shown to be very helpful to accurately quantify expression levels of lowly expressed genes and non-coding genes [18]. To evaluate our diagnostic algorithm, we developed a testing scenario based on simulated patients presenting with all clinical features listed in OMIM for a certain disease or syndrome. This validation test showed that for 49% of the diseases the causative gene ranks in the top 5%. We also investigated the OMIM “provisional” category of genes for which there is limited evidence. Both the OMIM disease-gene annotation and the provisional annotations perform significantly better than a random permutation. While we do find a small but significant difference in prediction performance between the provisionally annotated genes and the more established disease associated genes, we conclude, based on our findings, that these provisional OMIM annotations are generally of similar reliability to the other OMIM disease annotations.

Benchmarking on sequence data of patients with a known genetic diagnosis revealed that GADO returned the real causative variant within the top 3 results for 41% of the samples, indicating the potential power of GADO for a large number of diseases. Finally, in seven patients, GADO was able to identify potential novel disease genes that are strong candidates based on literature or functional evidence. For other cases we have identified genes with a strong prediction score harboring variants that might explain the phenotype. However, since very little is known about these genes it is not yet possible to draw firm conclusions. Hopefully this will become possible in the near future through initiatives like Genematcher [40].

### Potential to discover novel human disease genes

Over the last decade, several computational tools have been developed to prioritize variants in genes. Some, such as GAVIN, focus on variant filtering and prioritization based on deleteriousness scores, allele frequency and inheritance model [9]. Other methods measure the similarity between the clinical manifestations observed in a patient and those representing each of the diseases in a database or literature. Exomiser is closely related to GADO as it prioritizes genes based on specified HPO terms and also infers HPO annotation for unknown genes [14]. The gene prioritization by Exomiser is based on the effects of orthologs in model organisms and applies a guilt-by-association method using proteinprotein associations provided by STRING [41]. Exomiser performs better than GADO in ranking known disease-causing genes (supplementary figure 3, supplementary table 4) and is also able to identify potential new genes in human disease. However, Exomiser has a limitation in that only a subset of the protein-coding genes has orthologous genes in other species for which a knockout model also exists. Additionally, the used STRING interactions are biased towards well studied genes and rely heavily on existing annotations to biological pathways (supplementary figure 4). There are however, still 3,922 protein-coding genes that are not currently annotated in any of the databases we used, and there are even more non-coding genes for which the biological function or role in disease is unknown. Since GADO does not rely on prior knowledge, it can be used to prioritize variants in both coding *and* non-coding genes (for which no or limited information is available). GADO thus enables the discovery of novel human disease genes and can complement existing tools in analyzing the genomic data of patients who have a broad spectrum of phenotypic abnormalities.

### Limitations

The gene predictability score indicates for which genes we can reliably predict phenotypic associations and for which genes we cannot based on gene co-regulation. This score gives insight into which genes are expected to perform poorly in our prioritization. We found strong correlation between these gene predictability scores and the gene prioritization z-scores. Thus, genes with a high predictability score have more accurate HPO term predictions. However, since our predictions primarily rely on co-activation patterns that we identified from RNA-seq data, our method does not perform well for genes where gene-expression patterns are not informative of their function. This could, for instance, be the case for proteins relying heavily on post-translation modifications for regulation or genes for which different transcripts have distinct functions. This last limitation can potentially be overcome by predicting HPO-isoform associations by using transcript-based expression quantification.

Insufficient statistical power to obtain accurate predictions may be another explanation for the low predictability scores of certain genes. This may be true for genes that are poorly expressed or expressed in only a few of the available RNA-seq samples. The latter issue we expect to overcome in the near future as the availability of RNA-seq data in public repositories is rapidly increasing. Initiatives such as Recount enable easy analysis on these samples [42], allowing us to update our predictions in the future, thereby increasing our prediction accuracy.

For some genes we are unable to predict annotated disease associations despite having a high gene predictability scores. Some genes, such as *TTR,* simply act in a manner unique to a specific phenotype. Other genes, such as *TMPO,* turned out to be false positive disease associations. These examples show that our gene predictability score has the potential to flag genes acting in a unique manner as well as genes that might be incorrectly assigned to a certain disease or phenotype.

We noted that the median prediction performance of HPO terms is lower compared to the other gene sets databases used in our study, such as Reactome. This may be due to the fact that phenotypes can arise by disrupting multiple distinct biological pathways. For instance, DCMs can be caused by variants in sarcomeric protein genes, but also by variants in calcium/sodium handling genes or by transcription factor genes [43]. As our methodology makes guilt-by-association predictions based on whether genes are showing similar expression levels, the fact that multiple separately working processes are related to the same phenotype can reduce the accuracy of the predictions (although it is often still possible to use these predictions as the DCM HPO phenotype prediction performance AUC = 0.76).

### Complexity

Given that nearly 5% of patients with a Mendelian disease have another genetic disease [44], it is important to consider that multiple genes might each contribute to specific phenotypic effects. Clinically, it can be difficult to assess if a patient suffers from two inherited conditions, which may hinder variant interpretation based on HPO terms. We showed that GADO can disentangle the phenotypic features of two different diseases manifesting in one patient by correlating and subsequently clustering the profiles of HPO terms describing the patient’s phenotype. If the HPO terms observed for a patient do not correlate, it is more likely that they are caused by two different diseases. An early indication that this might be the case for a specific patient can simplify subsequent analysis because the geneticist or laboratory specialist performing the variant interpretation can take this in consideration. GADO also facilitates separate prioritizations on subsets of the phenotypic features.

## Conclusion

Connecting variants to disease is a complex multistep process. The early steps are usually highly automated, but the final most critical interpretations still rely on expert review and human interpretation. GADO is a novel approach that can aid users in prioritizing genes using patient-specific HPO terms, thereby speeding-up the diagnostic process. It prioritizes variants in coding *and* non-coding genes, including genes for which there is no current knowledge about their function and those that have not been annotated in any ontology database. This gene prioritization is based on co-regulation of genes identified by analyzing 31,499 publicly available RNA-seq samples. Therefore, in contrast to many other existing prioritization tools, GADO has the capacity to identify novel genes involved in human disease. By providing a statistical measure of the significance of the ranked candidate variants, GADO can provide an indication for which genes its predictions are reliable. GADO can also detect phenotypes that do not cluster together, which can alert users to the possible presence of a second genetic disorder and facilitate the diagnostic process in patients with multiple non-specific phenotypic features. GADO can easily be combined with any filtering tool to prioritize variants within WES or WGS data and can also be used in gene panels such as PanelApp [45]. GADO is freely available at www.genenetwork.nl to help guide the differential diagnostic process in medical genetics.

## Materials and Methods

### Gene co-regulation and function predictions

We used publicly available RNA-seq samples from the European Nucleotide Archive (ENA) database [46] to predict gene functions and gene-HPO term associations. After processing and quality control we included 31,499 sample for which we have expression quantification on 56,435 genes (supplementary methods 1). We performed a PCA on the gene correlation matrix and selected 1,588 reliable principal components (PCs) (Cronbach’s Alpha ≥ 0.7). We used the eigenvectors of these 1,588 PCs to predict gene functions and to predict HPO term associations [19]. We applied this methodology to the gene sets described by terms in the following databases: Reactome and KEGG pathways, Gene Ontology (GO) molecular function, GO biological process and GO cellular component terms and finally to HPO terms. We excluded terms for which fewer than 10 genes are annotated because predictions for smaller groups of genes are less accurate and might be misleading. Predictions were made for 8,657 gene sets in total.

The following steps were taken to obtain the gene prediction scores per gene set (**Figure** 1). First, for each PC, a student’s T-test was conducted between the eigencoefficients of the genes annotated to a particular gene set and a group of genes serving as a background. This background consisted of the genes annotated to any term in a specific database, excluding those annotated to the current term. Second, the resulting p-values of the T-test were transformed into a z-score, which indicate to which extend each PC represents a part of the biology underlying a gene set. This is done for each PC, resulting in a profile how important each PC is for a gene set. Finally, to predict which genes can be associated to a particular gene set, we correlated the 1,588 T-test z-scores for that gene set (as calculated above) with the 1,588 eigenvector coefficients of a gene. The p-value of this correlation indicates the fit between a gene and a pathway / HPO term, these p-values were transformed to predictions z-scores. When a gene was already explicitly annotated to a gene-set and we wanted to predict whether that gene is involved in that gene set, then there is a small circular bias as the predictions profile of this set was partly calculated based on this gene. To remove this bias, the 1,588 z-scores for a gene set were first re-calculated while assuming this gene is not involved in that gene set, after which the gene prediction was made.

To determine the accuracy of our predictions we assessed our ability to predict back known gene set annotations. For each gene-set, we calculated an Area Under the Curve (AUC), using a Mann-Whitney U test, on the predictions z-scores of the genes that are part of a set versus those that are not part of a set. These AUCs indicate how accurate the predictions were, with an AUC of 1 indicating perfect predictions and an AUC of 0.5 indicating no predictive power. The average AUC for each category was calculated based on all gene sets with at least 10 annotated genes and with a p-value ≤ 0.05 (Bonferroni corrected for the number of pathways in a database).

### Gene predictability scores

To explain why for some genes we cannot predict known HPO annotation, we have established a gene predictability score. We have calculated this gene predictability using the prioritization z-scores based on Reactome, GO and KEGG. For each gene and for each database we calculated the skewness in the distribution of the prioritization z-scores of the gene sets. We used the average skewness as the gene predictability score.

### GADO predictions

To identify potential causative variants in patients, we used HPO terms to describe a patient’s features. We only used the HPO terms which have significant predictive power (based on the p-value of U test to calculate the AUC). If the predictions for a patient’s HPO term were not significant, the parent/umbrella HPO terms were used (supplementary figure 1). The online GADO tool suggests the parent terms from which the user can then select which terms should be used in the analysis. The gene prediction z-scores for an HPO term were used to rank the genes. If a patient’s phenotype was described by more than one HPO term, a meta-analysis was conducted. In these cases a weighted z-score was calculated by adding the z-scores for each of the patient’s HPO terms and then dividing by the square root of the number of HPO terms [24]. The genes with the highest combined z-scores are predicted to most likely candidate causative genes for a patient. This analysis can be conducted at: https://www.genenetwork.nl.

### Validation of disease-gene predictions

To benchmark our method we used the OMIM morbid map [5] downloaded on March 26, 2018, containing all disease-gene-phenotype entries. From this list, we extracted the disease-gene associations, excluding non-disease and susceptibility entries. We extracted the provisional disease-gene associations separately. For each disease in OMIM, we used GADO to determine the rank of the causative gene among all genes in the OMIM morbid map. For this we used all phenotypes annotated to the OMIM disease. If any of the HPO terms did not have significant predictive power, the parent terms were used.

To determine if these distributions were significantly different from what we expect by chance, we permuted the data. We replaced the existing gene-OMIM annotation but assigned every gene to a new disease (keeping the phenotypic features for a disease together), assuring that the randomly selected gene was not already annotated to any of the phenotypes of the original gene.

### Cohort of previously solved cases

To test if GADO could help prioritize genes that contain the causative variant, we used 83 samples of patients who were previously genetically diagnosed through whole exome analysis or gene panel analysis. These samples encompass a wide variety of different Mendelian disorders (supplementary table 2). To assess which genes harbor potentially causative variants, we first called and annotated the variants from the exome sequencing files (Supplementary methods 3). For 11 of the previously solved cases, GAVIN did not flag the causative variant as a candidate. To be able to include these samples in our GADO benchmark, we added the causative genes for these cases manually to the candidate list.

The phenotypic features of a patient were translated into HPO terms, which were used as input to GADO. Here we only used features reported in the medical records prior to the molecular diagnosis. If any of the HPO terms did not have significant predictive power, the parent terms were used. From the resulting list of ranked genes, the known disease genes harboring a potentially causative variant were selected. Next, we determined the rank of the gene with the known causative variant among the selected genes. If a patient harbored multiple causative variants in different genes, in case of di-genic inheritance or two inherited conditions, the median rank of these genes was reported (supplementary table 2).

### Unsolved cases cohorts

In addition to the patients with a known genetic diagnosis, we tested 38 unsolved cases (supplementary table 3). These are patients with mainly cardiomyopathies or developmental delay. All patients were previously investigated using exome sequencing, by analyzing a gene panel appropriate for their phenotype. To allow discovery of potential novel disease genes, we used GADO to rank genes with candidate variants (Supplementary methods 3). For genes with a prediction z-score ≥ 5, a literature search for supporting evidence was performed to assess whether these genes are likely candidate genes.

### Website

To make our method and data available we have developed a website available at www.genenetwork.nl that can be used to run GADO, lookup gene functions predictions, visualize networks using co-regulations scores and perform function enrichments of sets of genes (Supplementary methods 4).

## Description of Supplemental Data

Supplementary methods 1. Processing and quality control of public RNA-seq data

Supplementary methods 2. Benchmark comparison with Exomiser

Supplementary methods 3. Variant calling and processing of benchmark samples

Supplementary methods 4. GeneNetwork website

Supplementary figure 1. Selection of parent HPO term if GADO does not have significant predictive power for query term

Supplementary figure 2. Comparison of GADO performance with the level of evidence for each cardiomyopathy-related gene

Supplementary figure 3. Comparison between GADO and Exomiser rankings

Supplementary figure 4. Correcting for biases in co-expression networks

Supplementary figure 5. Histogram of the gene types included in our analyses

Supplementary figure 6. PCA plot of 36,761 samples

Supplementary figure 7. Investigation of principal components capturing technical biases

Supplementary figure 8. Variance explained by first 1588 PCs

Supplementary figure 9. Visualization of PC1 to PC 10 of PCA over gene correlation matrix

Supplementary figure 10. Outlier genes in PC 8 and PC 9 of PCA over gene correlation matrix

Supplementary figure 11. PC sample scores to distinguish different tissues

Supplementary figure 12. Outlier samples in PC sample scores of PC 8 and PC 9

Supplementary table 1. A list of samples annotated in the European Nucleotide Archive June 30, 2016

Supplementary table 2. A list of 83 diagnosed patients with Mendelian disorders and corresponding predictions with GADO

Supplementary table 3. A list of 38 undiagnosed patients with suspected Mendelian disorders

Supplementary table 4. A comparison between GADO and Exomiser predictions using a list of 83 diagnosed patients with Mendelian disorders

## Acknowledgments

We are grateful for the participation of the patients and their parents in this study. We thank Kate Mc Intyre for editing the manuscript and Marieke Bijlsma, Gerben van der Vries, Sido Haakma and Pieter Neerincx for support with the computational analyses. This work was carried out on the Groningen Center for Information Technology (Strikwerda, W. Albers, R. Teeninga, H. Gankema and H. Wind) and Target storage (E. Valentyn and R. Williams). Target is supported by Samenwerkingsverband Noord Nederland, the European Fund for Regional Development, the Dutch Ministry of Economic Affairs, Pieken in de Delta and the provinces of Groningen and Drenthe. Wouter P. te Rijdt is supported by Young Talent Program (CVON PREDICT) grant 2017T001 from the Dutch Heart Foundation. Netherlands Heart Institute, Utrecht, the Netherlands. This work is supported by a grant from the European Research Counsil (ERC Starting Grant agreement number 637640 ImmRisk) to Lude Franke and a VIDI grant (917.14.374) from the Netherlands Organisation for Scientific Research (NWO) to Lude Franke. This work was supported by BBMRI-NL, a research infrastructure financed by the Dutch government (NWO 184.021.007).

